# High yield preparation of outer-membrane protein efflux pumps by *in vitro* refolding is concentration dependent

**DOI:** 10.1101/2020.09.14.296756

**Authors:** S. Jimmy Budiardjo, Ayotunde Paul Ikujuni, Emre Firlar, Andrés Cordova, Jason T. Kaelber, Joanna S.G. Slusky

## Abstract

Overexpression of tripartite efflux pump systems in gram-negative bacteria are a principal component of antibiotic resistance. High-yield purification of the outer membrane component of these systems will enable biochemical and structural interrogation of their mechanisms of action and allow testing of compounds that target them. However, preparation of these proteins is typically hampered by low yields requiring laborious large-scale efforts. If refolding conditions can be found, refolding these proteins from inclusion bodies can lead to increased yields as compared to membrane isolations. Here, we develop a concentration-dependent folding protocol for refolding TolC, the outer membrane component of the antibiotic efflux pump from *Escherichia coli*. We show that by our method of re-folding, homotrimeric TolC remains folded in SDS-PAGE, retains binding to an endogenous ligand, and recapitulates the known crystal structure by single particle cryoEM analysis. We find that a key factor in successful re-folding is a concentration dependence of TolC oligomerization. We extended the scheme to CmeC, a homologous protein from *Campylobacter jejuni*, and find that concentration-dependent oligomerization is a general feature of these systems. Because outer-membrane efflux pump components are ubiquitous across gram-negative species, we anticipate that incorporating a concentration step in re-folding protocols will promote correct refolding allowing for reliable, high-yield preparation of this family of proteins.

## Introduction

Gram-negative bacteria assemble tripartite efflux pump machineries that are responsible for extruding toxic compounds such as heavy metals and antibiotics (recently reviewed by (Neuberger et al., 2018)). These systems have been found to be overexpressed in multidrug resistant clinical isolates (Swick et al., 2011). Tripartite efflux pump systems form intermembrane complexes between the inner and outer membranes made up of three components (Daury et al., 2016). The inner membrane component is a pump driven by an energy-dependent mechanism such as proton motive force or ATP hydrolysis that extrudes the substrates into the extracellular environment. The outer membrane component of these pumps forms the channel through the outer membrane and into the extracellular environment. The inner and outer membrane components are linked by a periplasmic adapter protein. Structural and biochemical characterization of these tripartite efflux pumps are necessary in order to better understand antibiotic resistance mechanisms and ways to combat them.

An inherent limitation when working with membrane proteins is generating large enough quantities for experiments. For expression of outer membrane proteins (OMPs), it is typical to utilize expression culture volumes between 10-20L (Noinaj et al., 2016). Even if using a bioreactor, processing such large volumes is laborious and time consuming. Poor yields are in part due to limited space on the membrane to accommodate overexpressed protein compared to the cytoplasm. Heterologous expression of proteins from different species in a host organism like *Escherichia coli* (*E. coli*) can be hindered by differences of lipid composition or toxicity of the native protein in the membrane which can also reduce yield (Bannwarth and Schulz, 2003). It has long been known that poor yield can be overcome by cytoplasmic accumulation into inclusion bodies whereby 20% of *E. coli* cellular volume is occupied by recombinant protein (Williams et al., 1982). By deleting the N-terminal signal sequence that directs the polypeptide chain across the inner membrane into the periplasm, OMPs accumulate in the cytoplasm as inclusion bodies (Schmid et al., 1996). However, successful over expression is just one requirement. Screening conditions for solubilizing inclusion bodies in a denaturant followed by refolding the protein in a lipid mimetic environment into its native state is not a trivial task.

Successful refolding for many OMPs into detergent micelles or lipid vesicles has been documented (Surrey and Jahnig, 1992, Schmid et al., 1996, Visudtiphole et al., 2005, Burgess et al., 2008). Typically, these OMPs form the prototypical enclosed barrel comprised of a single polypeptide chain of 8-24 stranded β-sheets. In contrast, outer membrane channels of efflux pump systems such as TolC from *E. coli* are different from most other outer membrane proteins (Franklin et al., 2018). Specifically, these outer membrane barrels are comprised of three separate polypeptide chains that come together to form a single β-barrel (**Figure 1A**). The polymeric protein topology adds an additional layer of complexity for correct assembly. Moreover, each chain contains a large soluble α-helical barrel that extends beyond the membrane into the periplasm. Therefore, refolding conditions require refolding of an amphipathic protein where half of the protein is soluble while the other half is membrane embedded.

**Figure 1:**
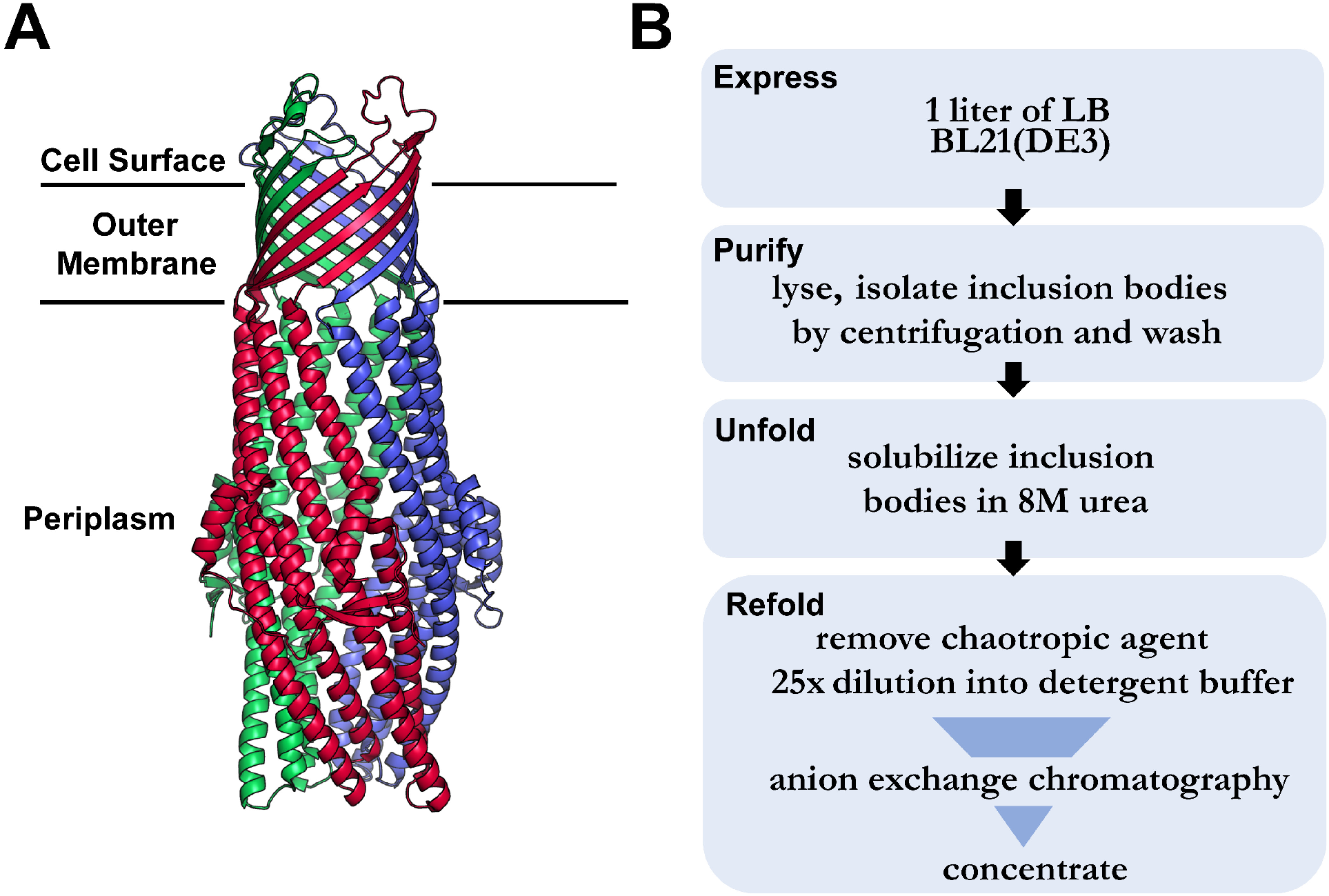
How to refold TolC *in vitro*. (A) The folded structure of outer membrane protein TolC from *Escherichia coli* (from PDB 1EK9 (Koronakis et al., 2000)). TolC is a homotrimer. Each color (red, green and blue) is a different chain. (B) TolC expression, purification and refolding scheme

To our knowledge, there have only been three examples of successful refolding of outer membrane channels of efflux pump systems: OprM from *Pseudomonas aeruginosa* (Charbonnier et al., 2001), AggA from *Shewanella oneidensis* (Theunissen et al., 2009), and Slr1270 from *Synechocystis* 6803, a cyanobacteria (Agarwal et al., 2014). Here we describe the expression, purification, and refolding of the efflux pump protein TolC from *E. coli*, the most well studied of the tripartite efflux pumps. Despite being the prototypical membrane channel for tripartite efflux pump systems, no reports have been published on refolding of TolC and all structural and functional studies have required laborious membrane purification (Koronakis et al., 1997, Koronakis et al., 2000, Tikhonova and Zgurskaya, 2004, Zakharov et al., 2004, Tikhonova et al., 2007, Tikhonova et al., 2009, Daury et al., 2016, Zakharov et al., 2016). As a test case for the generalizability of our refolding protocol, we extended this scheme to CmeC, a homologous protein from *Campylobacter jejuni*. We find that a critical step in the successful refolding of outer membrane channels of efflux pump systems is the concentration of the protein sample after removal of the chaotropic agent to transition the monomeric species to the fully folded trimer. We believe that including this step into refolding protocols will aid in the successful refolding of other outer membrane efflux pump proteins.

## Results

The general scheme for refolding TolC is outlined in (**Figure 1B**). After solubilization of the inclusion bodies in 8M urea, a 25-fold dilution of TolC is performed into a buffer containing *n*-octylpolyoxyethylene detergent micelles to reduce the chaotropic agent below a concentration at which it can effectively keep the protein unfolded. This initiates refolding. After rapid dilution, the protein is applied to an anion exchange column to both recover the protein and to remove the remaining 0.32 M urea. After the protein is eluted it is concentrated to oligomerize TolC.

To assess proper refolding, we performed a heat modifiability assay in which folding can be determined by a shift in apparent molecular weight between the folded and unfolded states using semi-native SDS-PAGE. Folded OMPs exhibit resistance to SDS unfolding and thus a natively-folded OMP is more compact and will migrate faster than an unfolded OMP (after heating at 95 °C) (Rosenbusch, 1974, Nakamura and Mizushima, 1976, Heller, 1978, Noinaj et al., 2015). Additionally, because TolC exists as a trimer there will be a large molecular weight difference between properly folded trimers (154,364 Da) and monomers (51,454 Da). After anion exchange chromatography, the protein remains monomeric as there is no molecular weight shift if the protein is boiled (fully unfolded) or if it is left unboiled (**Figure 2, left**). However, if the elution fraction from anion exchange is concentrated the monomeric state transitions into the trimeric state (**Figure 2, right**). We find that full trimerization is achieved at 50 μM and above. As seen with other OMPs, the trimeric state migrates faster at ~100 kDa versus its calculated 154 kDa molecular weight since it is presumably more compact than an extended polypeptide of the same length.

**Figure 2:**
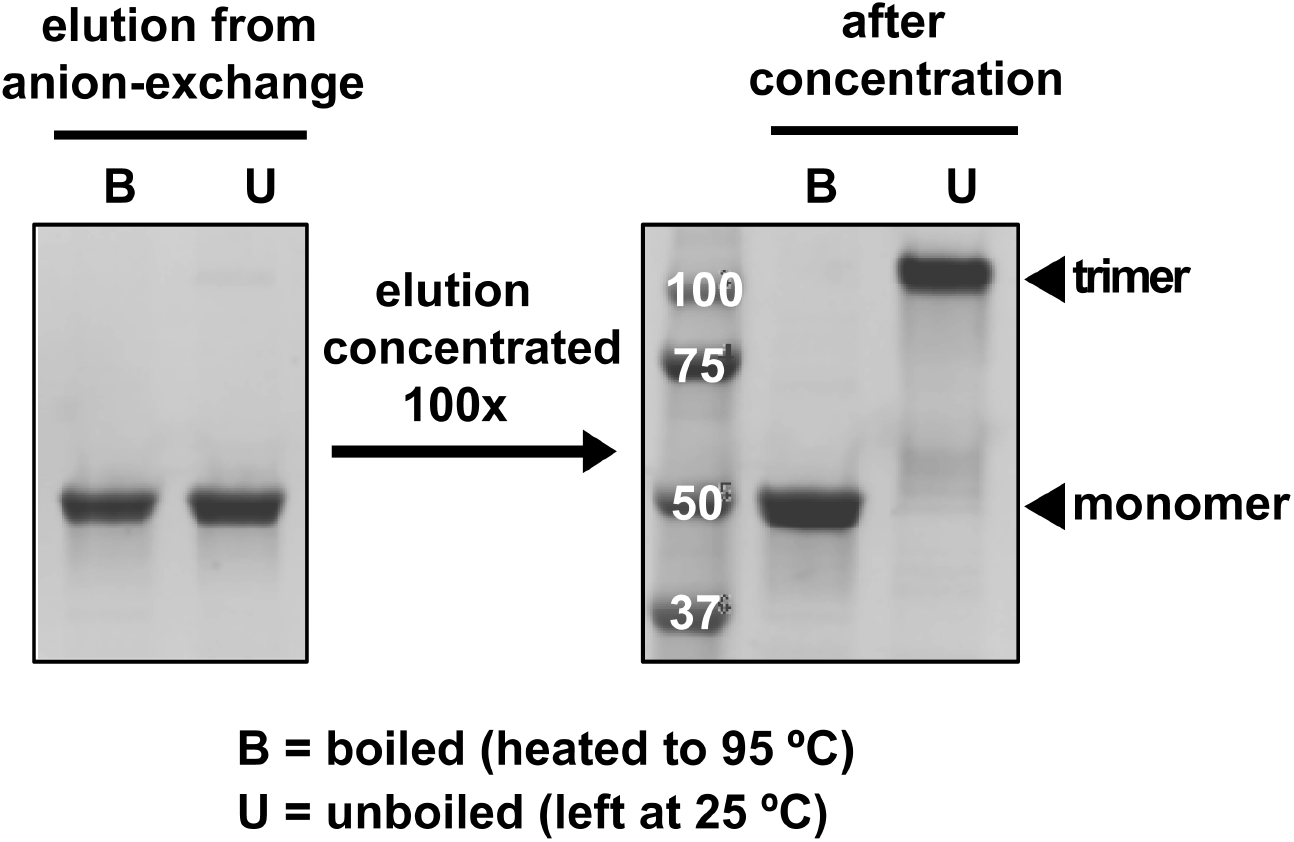
Refolding of TolC is concentration dependent. Heat modifiability assay to assess folding of TolC indicates that TolC transitions into trimers upon concentration.

To determine if TolC folded correctly, we assessed whether its binding interface remains intact and able to bind to an endogenous protein ligand. Colicin 5 (Col5) is a bacteriocin secreted by *E. coli* to kill neighboring cells that hijacks surface exposed TolC to enter the cell (Pilsl and Braun, 1995). We expressed and purified residues 1-201 of Col5 and assessed TolC binding through co-elution on a Superdex size exclusion chromatography column. TolC (purple line) and Col5 (cyan line) alone elute at 10.6 mL and 13.7 mL, respectively (**Figure 3, left**). If TolC and Col5 are premixed before loading onto the column, the TolC peak shifts to 10.2 mL (black line) indicative of complex formation with Col5 forming a larger molecular weight species that elutes earlier. We then took the peak fraction from 10.2 mL (red arrow) and found that both proteins are present in the peak by SDS-PAGE analysis (**Figure 3, right**).

**Figure 3:**
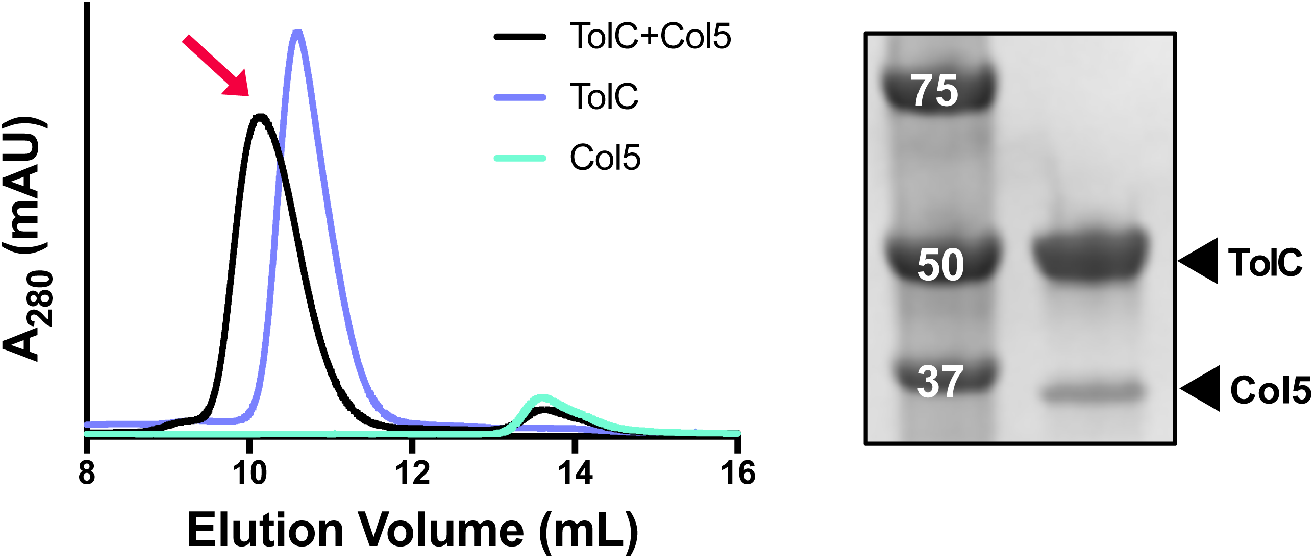
Refolded TolC binds to Col5. SEC chromatograms of TolC (purple), Col5 (cyan) and TolC+Col5 (black). TolC and Col5 were mixed at a 1:2 molar ratio 10ìM TolC:20ìM Col5. Peak fraction from 10.2 mL (red arrow) used for SDS-PAGE analysis. Sample was boiled as reflected by only monomeric TolC species.

To fully confirm proper refolding, we sought to determine TolC quaternary structure by single particle cryoEM. To avoid difficulties introduced by free detergent micelles in solution that could complicate single particle analysis, TolC was refolded using our new protocol and embedded into lipid nanodiscs. The “tube” shape of TolC was readily visible in raw micrographs, particularly when the ice was thinner than 30 nm (**Figure 4A**). Additionally, the “donut” shape is apparent where TolC molecules are oriented in the ice orthogonal to the side views and one is able to visualize the axis down barrel. Representative 2D classes for TolC in random orientations show strikingly similar features of the known solved crystal structure (Koronakis et al., 2000) where both the alpha helical periplasmic region and beta barrel crown wrapped by the nanodisc are resolved (**Figure 4B**). *Ab initio* reconstruction confirmed that TolC was trimeric, fully-folded, and structurally intact (**Figure 4C**). The Fourier shell correlation between the *ab initio* cryoEM reconstruction and the crystal structure of TolC (Koronakis et al., 2000) crossed 0.5 at 8Å, comparable to the resolution of the map. The residues which were ordered in the crystal structure were enclosed in the cryoEM density, while the disordered terminal residues were not observed; no significant deviations between the nanodisc-embedded cryoEM structure of TolC and previously-crystallized TolC were seen except that, in cryoEM, density was observed for the nanodisc in which the TolC trimer sits. Our *ab initio* reconstruction does not incorporate any previous information from solved structures of TolC and is fully data-derived, yet it recapitulates the crystal structures of TolC. We find that secondary structural elements of the protein are fully folded up to the last ordered residues in previous crystal structures.

**Figure 4:**
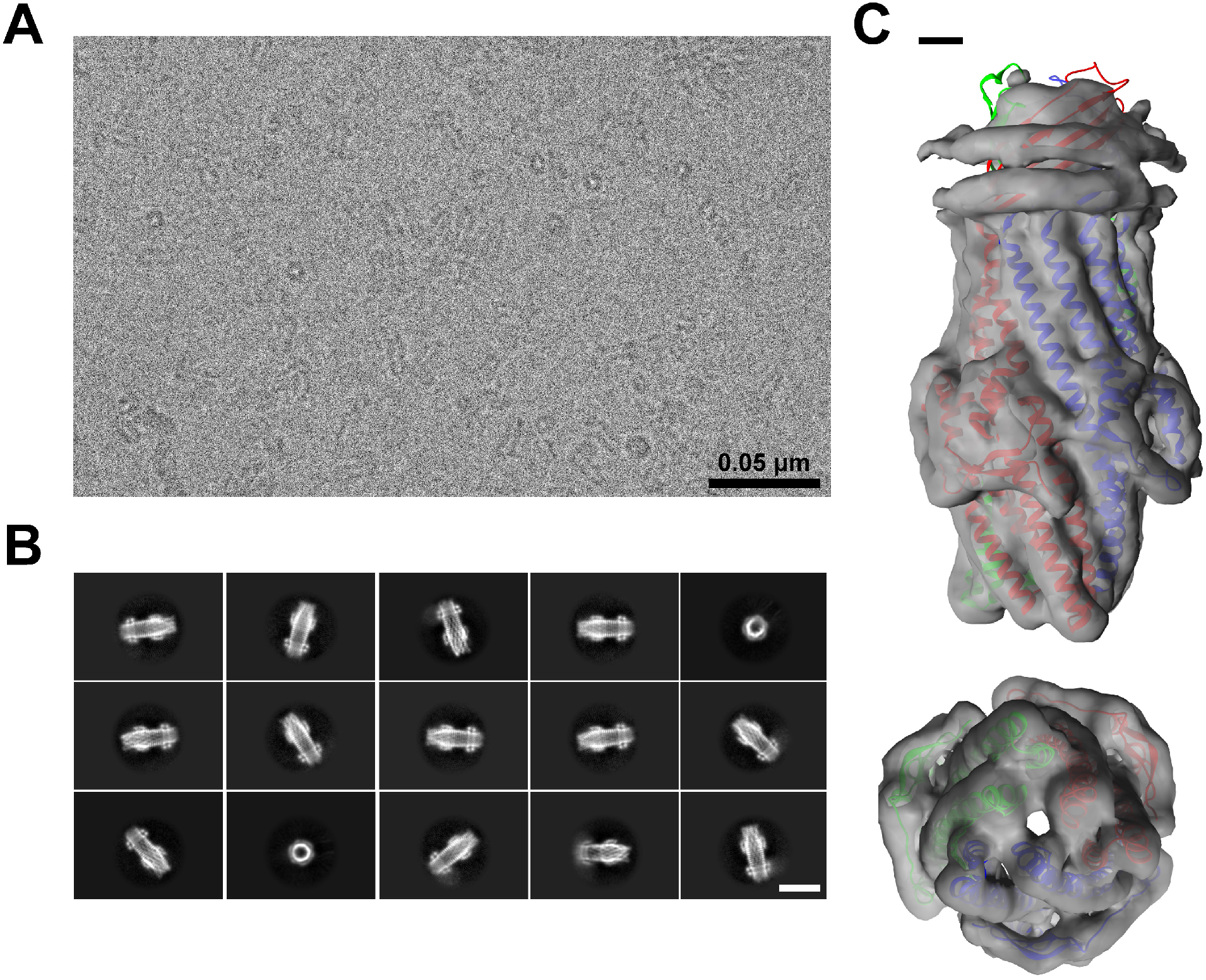
Single particle cryoEM reveals properly refolded TolC. (A) Representative micrograph of nanodisc embedded TolC with a scale bar in black representing 0.05 μm (B) Two-dimensional (2D) classes of nanodisc embedded TolC in side and top views with white scale bar representing 0.01 μm (C) *Ab initio* reconstruction of nanodisc-embedded TolC (translucent gray), top and side view, with TolC crystal structure (PDB: 1EK9, (Koronakis et al., 2000)) rigidly docked into the density and colored by chain. Scale bar 1 nm

We next sought to determine if concentration dependent refolding is a general feature of gram-negative polymeric outer membrane efflux pump channels. We applied the same protocol for refolding to CmeC, a homologous protein from the pathogenic species *Campylobacter jejuni* in which refolding has not been previously reported. CmeC is structurally homologous to TolC (Wu et al., 2014) (**Figure 5A**) but only has 24% sequence similarity as measured by BlastP (Altschul et al., 1997, Altschul et al., 2005). As seen with TolC, after rapid dilution and anion exchange chromatography CmeC remains monomeric (**Figure 5B, left**). Upon concentration a second higher molecular weight species appears at ~80 kDa (**Figure 5B, right**). Although this behavior is consistent with TolC, the molecular weight is ~20 kDa smaller than the known TolC trimer as analyzed by SDS-PAGE. We next compared the elution volume of CmeC to TolC by size exclusion chromatography and found that CmeC elutes at 10.6 mL as a single peak identical to TolC (**Figure 5C**). This suggests that CmeC is fully refolded and is the same molecular weight in solution as TolC. The smaller size of the trimeric species is likely explained by the phenomena of the folded species being more compact as it migrates through the gel and is more pronounced in CmeC versus TolC. Additionally, because the conditions of size exclusion chromatography are more native-like than semi-native SDS-PAGE, the fact there is a single peak indicates that CmeC is trimeric in solution and is less resistant to SDS mediated dissociation than TolC on the gel. To further confirm trimerization we also determined the molecular weight of CmeC by Matrix-assisted laser desorption/ionization (MALDI) mass spectrometry of crosslinked CmeC. Amine-reactive disuccinimidyl suberate (DSS) was used to crosslink concentrated CmeC to covalently trap the trimeric species. MALDI analysis of both the crosslinked and un-crosslinked shows a broad peak of the trimer (~164 kDa) in only the sample treated with DSS (**Figure 5D**). The low peak intensity is typical of larger proteins that do not fly well in MALDI preparations. The broad peak is due to heterogenous crosslinking combinations by DSS of 35 lysines per monomer of CmeC. The mass + 2 × cation aduct (~80 kDa) of the trimer is also only visible in the sample treated with DSS. The CmeC monomer (~50 kDa) is visible in both the DSS crosslinked and un-crosslinked sample (as is the monomer 2 × mass + cation aduct, ~100 kDa).

**Figure 5:**
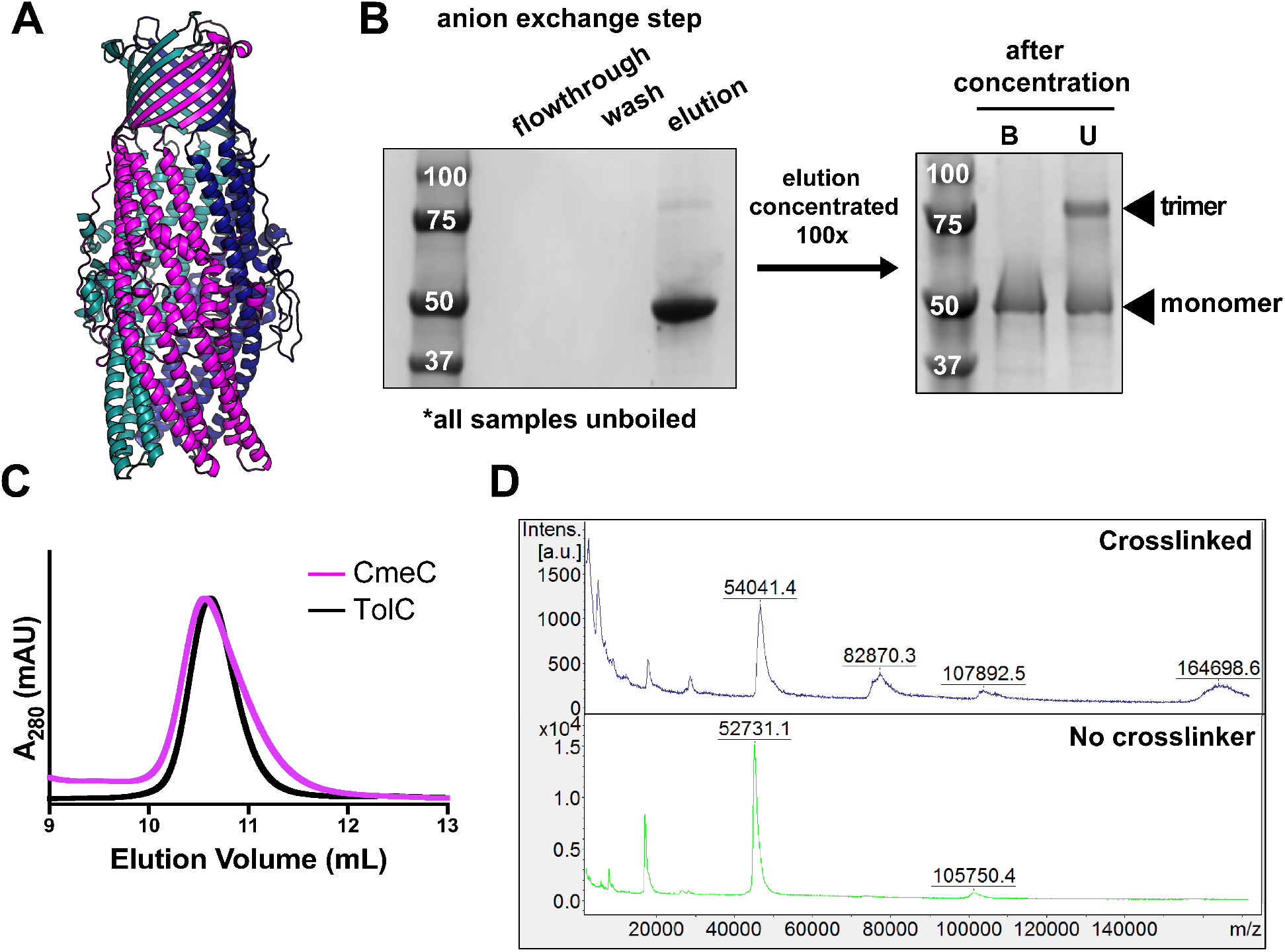
Refolding of CmeC is concentration dependent. (A) The folded structure of outer membrane protein CmeC from *Campylobacter jejuni* (from PDB 4MT4). CmeC is a homotrimer. Each color (pink, teal, and navy blue) is a different chain. (B) Heat modifiability assay to assess folding of CmeC indicates that CmeC transitions into trimers upon concentration like TolC. (C) Size exclusion chromatography of CmeC (pink) versus TolC (black) (D) MALDI Mass Spectrometry of CmeC with (blue, top) and without (green bottom) covalent crosslinker disuccinimidyl suberate (DSS).

## Discussion

We have successfully refolded TolC from inclusion bodies by developing a protocol that involves a rapid dilution step followed by anion exchange chromatography and induction of trimerization by concentration. This method yields ~10 mg of trimeric TolC from 1L of expression culture which is a 5- to 10-fold higher yield than previous reports of isolating folded TolC from its native environment in the outer membrane (Koronakis et al., 1997). In addition to the increased yield, this protocol is less laborious than native membrane isolation. Single particle cryoEM revealed that TolC folded correctly and adopts the same characteristics of the known crystal structure. We have also shown that there is a concentration dependence on trimerization. We were able to apply the same refolding protocol to another trimeric efflux pump protein CmeC which adopts a strikingly similar fold with only 24% sequence similarity as measured by BlastP (Altschul et al., 1997, Altschul et al., 2005).

Of particular significance is the concentration dependent trimerization of TolC and CmeC. OprM and AggA were both folded in similar schemes (Charbonnier et al., 2001, Theunissen et al., 2009) in which inclusion bodies were solubilized in either 6M guanidine hydrochloride or 8M urea followed by a 20- to 25-fold dilution into a buffer containing detergent. To recover the sample, they were loaded onto an anion exchange followed by a nickel column for OprM or just a nickel affinity column for AggA. Both TolC and CmeC were fully monomeric, even with no chaotropic agent after anion exchange. In principle, using protein concentrations above the trimerization inducing threshold may result in trimers in earlier steps as recovering the large volumes from rapid dilution on a column is effectively concentrating the sample. However, use of high concentrations is typically disfavored in steps like rapid dilution as it can promote aggregation resulting in reduced yield. In the case of AggA, the sample was concentrated after nickel column elution to be further purified by gel filtration. Slr1270 was refolded using a stepwise decrease in urea by dialysis (Agarwal et al., 2014) followed by concentration for gel filtration. Checking the oligomerization states of OprM, AggA and Slr1270 at each step may show that these protocols implicitly captured the concentration dependence of trimerization.

Our experiments on TolC and CmeC potentially deconvolute a possible multistep process in folding. Since there is no urea after the anion exchange step, yet all the protein remains monomeric, it is compelling to consider that below a concentration threshold, the protein exists as a pre-folded monomeric intermediate (**Figure 6, top**). In this scenario, rapid dilution initiates refolding followed by anion exchange chromatography that harvests the protein from a large volume while also removing residual urea leading to folded monomers. Once folded monomers are in solution, they are primed to form oligomers once a concentration threshold has been passed. However, without secondary structure analysis of the monomeric species we cannot discount the possibility of unfolded monomers that form secondary structure upon oligomerization (**Figure 6, bottom**). It is currently unknown how these proteins fold and insert into the outer membrane in cells so further investigation into pre-folded intermediate states may shed light on how the process occurs *in vivo*.

**Figure 6:**
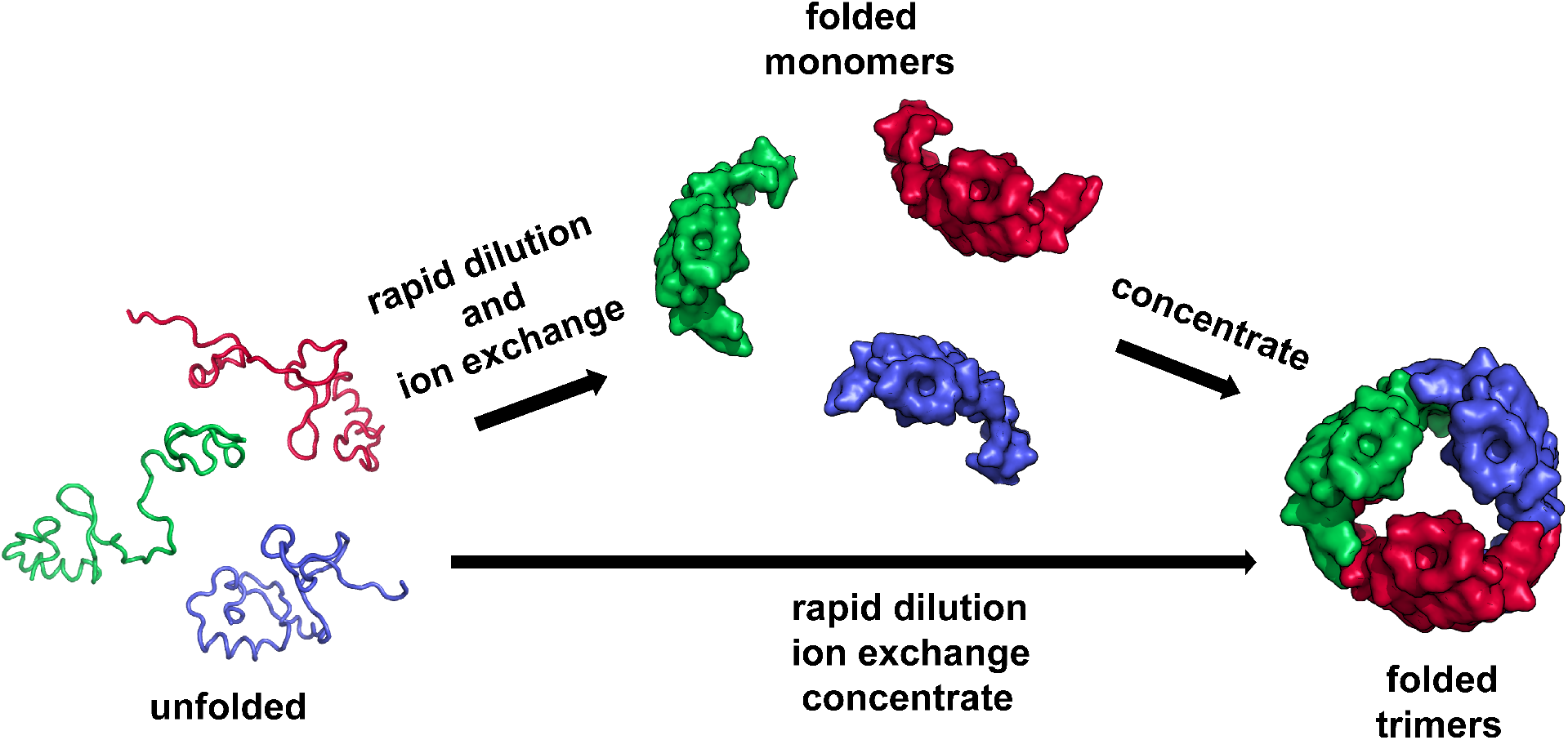
Potential pathways of *in vitro* TolC refolding

Efflux pump systems like these are universal in gram-negative bacteria. *Pseudomonas aeruginosa* alone encodes three such outer membrane efflux pump proteins within its genome: OprM, OprJ and OprN (Horna et al., 2018). We believe that the findings reported here will be broadly applicable to other homologs of TolC although the choice of chaotropic agent used to solubilize inclusion bodies, detergent choice for refolding, chromatography resins will likely be protein dependent. This will enable *in vitro* refolding with high yield for biochemical and structural studies.

## Materials and Methods

### *E. coli* strains

BL21(DE3) were used for expression of the Col5 and TolC into inclusion bodies.

### Protein Expression and Purification

#### TolC and CmeC (inclusion body isolation)

To express TolC as inclusion bodies, the signal sequence was deleted using the forward primer 5’-CATGGTCTGTTTCCTGTGTGAAATTG -3’ and reverse primer 5’-GAGAACCTGATGCAAGTTTATCAGC -3’. The gene for CmeC was synthesized (GenScript) and cloned into pTrc99a. Sequence confirmed TolC and CmeC in pTrc99a was used to transform E. coli BL21 (DE3) cells and plated in a LB agar plate 100 μg/mL carbenicillin. A single colony from the plate was used to inoculate a 20 mL LB media containing 100 μg/mL carbenicillin and incubate at 37 °C, 250 r.p.m. overnight (~ 16 hours). The starter culture was used to inoculate 1L of LB media containing 100 μg/mL carbenicillin at 37 °C, 250 r.p.m. until the optical density at 600 nm reached ~0.6. The expression of TolC was induced with IPTG to a final concentration of 1 mM and incubated at 37 °C with shaking at 250 r.p.m. for 4 hours. Cells were harvested by centrifugation at 3,739 g for 25 minutes at 4 °C. Cell pellets were resuspended in Lysis buffer (TBS, 5 mM MgCl_2_, 0.2 mg/mL lysozyme, 5 μg/mL DNase, 1mM PMSF, 1% (v/v) Triton X-100) at 3 mL/1g of cell pellet and lysed via sonication (4 minutes, 2s on, 8s off, 40% amplitude, QSonica Q500 with 12.7 mm probe) in an ice bath. The lysate was centrifuged at 3,739 g for 25 minutes at 4 °C and the inclusion body pellet was resuspended in 40 mL wash buffer (20 mM Tris pH 8.0, 100 mM NaCl, 1 mM EDTA, 1% (v/v) Triton X-100). The lysate was sonicated in an ice bath for an additional 3 minutes and inclusion bodies were recovered by centrifugation at 3,739 g for 25 min at 4 °C. Inclusion body pellets were washed one final time by resuspending in wash buffer and centrifuged at 3,739 g at 4 °C for 25 min. The inclusion bodies pellets were then resuspended in storage buffer (20 mM Tris-HCl, pH.0 8) aliquoted and stored at −20 °C until further use.

### Refolding TolC and CmeC

#### Refolding by concentration

Inclusion bodies were thawed and centrifuged at 3,739 g for 25 min at 4 °C. The inclusion body pellet (~60 mg) was dissolved in urea buffer (20 mM Tris-HCl, 8 M urea pH 8) at a concentration ~6.8 mg/mL and incubated at 37 °C for 30 minutes. To initiate refolding, solubilized TolC or CmeC was diluted 25-fold into refolding buffer (20 mM Tris-HCl, 1% (w/v) n-octylpolyoxyethylene (C8POE), pH 8.0) on a stir plate at 100 r.p.m. The solution was passed through a 0.22 μm pore size vacuum filter and loaded onto a 5 mL HiTrap™ Q HP anion exchange column equilibrated with wash buffer (20 mM Tris-HCl pH 8.0, 1% (w/v) C8POE). The column was then washed with 5 column volumes of wash buffer. The protein was eluted with 5 column volumes of elution buffer (20 mM Tris-HCl pH 8.0, 1% (w/v) C8POE, 500 mM NaCl). TolC and CmeC remains monomeric after rapid dilution and Q column purification as determined by SDS-PAGE. Formation of trimeric TolC or CmeC was initiated by concentration of the elution fractions in an Amicon protein concentrator with a molecular weight cut off of 10 kDa by centrifugation at 4,000 x g at 4 °C until the volume has reduced by 100-fold. Formation of trimeric TolC or CmeC from monomers was confirmed using SDS-PAGE.

### Col5

The gene corresponding to residues 1-201 of colicin 5 (colE5-T) was synthesized (GenScript) and cloned into pET21(+) containing a C-terminal 6x histidine tag. Plasmids were transformed into BL21(DE3) and plated on LB + agar + 100 μg/mL carbenicillin. A single colony was inoculated into 20 mL of LB + 100 μg/mL carbenicillin and grown at 37 °C with shaking at 250 r.p.m. overnight. The next morning 1L TB broth + 10 mM MgCl2 + 100 μg/mL carbenicillin was inoculated with 20 mL of the overnight starter culture and grown at 37 °C with shaking at 250 r.p.m. until OD_600_ reached 1.0 and induced with 1 mM IPTG. The temperature was reduced to 15 °C with shaking at 250 r.p.m. and incubated for 24 hours. Cell pellets were harvested by centrifugation at 4,000 g for 30 minutes at 4 °C. Cell pellets were resuspended in lysis buffer (TBS, 5 mM MgCl_2_, 0.2 mg/mL lysozyme, 5 μg/mL DNase, 1mM PMSF, 20 mM imidazole) at 3 mL/gram of cell pellet. Cells were lysed by sonication in an iced water bath (3 minutes, 2s on, 8s off, 40% amplitude, QSonica Q500 with 12.7 mm probe). The cytoplasmic fraction was isolated by centrifugation at 50,400 g at 4 °C for 1 hour. The supernatant was filtered through 0.22 μm and applied to a 5 mL HisTrap™ FF column and purified using an ÄKTA FPLC system with a 20 column volume wash step with binding buffer (20 mM Tris, 400 NaCl, 25 mM imidazole) and eluted using a linear gradient from 0-100% elution buffer (20 mM Tris, 400 NaCl, 500 mM imidazole) in 10 column volumes. Colicin-containing fractions were pooled and concentrated to 2 mL and applied onto a HiLoad 16/60 Superdex 200 pg column and eluted with 1.5 column volumes in 20 mM Tris, 40 NaCl.

### Co-elution

TolC and colicin 5 were both buffer exchanged into co-elution buffer (20 mM Tris pH 8.0, 40 mM NaCl, 0.05% n-dodecyl-β-D-maltoside) using PD-10 desalting columns. Binding was determined by co-elution on a Superdex 200 Increase 10/300 SEC column. TolC and Col5 were mixed at a 1:2 molar ratio (5 μM TolC trimer to 10 μM Col5) and incubated at room temperature for 1 hour. The sample was eluted in 1.5 column volumes into 20 mM Tris pH 8.0, 40 mM NaCl, 0.05% n-dodecyl-β-D-maltoside and fractions were collected at every 0.5 mL.

### Embedding TolC into lipid nanodiscs

The plasmid for membrane scaffold protein was MSP1D1 in pET28a was purchased from Addgene and prepared as previously described (Hagn et al., 2018). Lipids (POPC) in chloroform (Avanti Polar Lipids) were dried under nitrogen and lyophilized to remove any residual chloroform. Lipids were solubilized in 20 mM Tris pH 8.0, 100 mM NaCl, 0.5 mM EDTA, 100 mM cholate to a final concentration of 50 mM. The tube was submerged in warm water and shaken until the solution cleared. The reaction mixture consisted of lipids, MSP1D1 and TolC mixed to a 36:1:0.4 ratio as previously described (Daury et al., 2016). Detergents were added to a final concentration of 0.1% n-dodecyl-β-maltoside and 20 mM cholate and the reaction incubated for 1 hour on ice. To initiate nanodisc formation, Bio-Beads SM-2 (Pre-washed in methanol and equilibrated with water) were added at 0.5 mg/mL of reaction and incubated at 4 °C on a rotary mixer overnight. The next morning the reaction was transferred into a tube with fresh Bio-Beads and incubated for 4 more hours. To separate TolC embedded nanodiscs from aggregates and empty nanodiscs, the sample was loaded onto a HiLoad 26/60 Superdex 200 pg column and eluted into 20 mM Tris pH 8.0, 40 mM NaCl. TolC containing fractions were concentrated to 4 mg/mL for cryoEM analysis.

### CryoEM

#### CryoEM sample preparation and data collection

UltrAuFoil R2/2 200 mesh TEM grids (Quantifoil Micro Tools) were prepared by 200 second glow discharge in 0.39 mBar of room air using a Pelco easiGlow (Ted Pella). 3 μl of TolC protein solution (4 mg/mL) was plunge frozen on these grids via Vitrobot Mark IV (FEI Company) using +5 force and 5 second blotting with grade 595 filter paper. Imaging was performed using a Talos Arctica (FEI Company) operated at 200kV with energy filtered K2 Summit (Gatan, Inc.) for detection. Images were acquired at a calibrated pixel size of 1.038Å/pixel in counting mode 33.04 e^−^/A^2^ total electron dose over 8 seconds (40 frames. Coma-compensated beam/image-shift automated data collection was performed in SerialEM (Mastronarde, 2018) using scripts of Chen Xu with minor modifications (available on request).

#### CryoEM data processing

16,433 micrographs were processed in CryoSPARC 2 (Punjani et al., 2017). Micrographs were imported as different image groups for the compensation of image shift dependent coma. Full frame motion correction was employed for the motion correction. CTFFIND 4.1.10 (Rohou and Grigorieff, 2015) was utilized to find the CTF parameters. 2,092,678 putative particles were extracted using template-based particle picking, with an amphipol-embedded TolC complex as the model (J.T. Kaelber, personal communication). After two rounds of 2D classification, 196,158 particles were retained for *ab initio* reconstruction to 8Å. No symmetry was imposed.

### Crosslinking and MALDI-TOF of CmeC

After concentration, CmeC was buffer exchanged into 100 mM sodium phosphate pH 7.4, 100 mM NaCl, 0.5% n-dodecyl-β-D-maltoside using a Bio-Spin 6 desalting column (Bio-Rad). CmeC (5 μM) was crosslinked with disuccinimidyl suberate (DSS) (500 μM) at room temperature for 5 minutes. The reaction was quenched with 5 mM Tris. Crosslinked CmeC was buffer exchanged into 20 mM Tris, 0.5 % Triton-X100 using a Bio-Spin 6 desalting column for mass spectrometry. CmeC was spotted onto the target plate in a sandwich of 1 μL of the matrix (sinapinic acid in 50% acetonitrile, 50% water and 0.1% trifluoroacetic acid) on the target plate which was allowed to dry, followed by 1 μL CmeC mixed 1:1 with matrix and allowed to dry and finally 1 μL of matrix. Mass spectra were collected on a Microflex LT (Bruker).

## Acknowledgments

We gratefully acknowledge Rik Dhar, Alex Little, and Jaden Anderson for discussions and feedback, Rajeev Misra for the pTrc vector containing the TolC gene and Vasileios Petrou for guidance on nanodiscs, Funding: The Gordon and Betty Moore Inventor Fellowship to JSGS and NIGMS award DP2GM128201 to Joanna Slusky and P20GM103638 and Kansas INBRE, P20 GM103418 to Jimmy Budiardjo.

We are also grateful for inspiration and mentorship from Stephen White. Our understanding of membrane protein folding is richer for your contributions and conferences are more fun when you are there. Thank you for leading the way in asking the important questions. Happy birthday and many happy returns.

